# A rapid ecological assessment for necrophagous flies (Diptera, Calyptratae) in a mosaic landscape of the Colombian Andes

**DOI:** 10.1101/2020.07.24.220491

**Authors:** Eduardo Amat, Claudia A. Medina

## Abstract

A necrophagous flies ensemble (Diptera, Calyptratae) was rapidly assessed in four coverages of an anthropogenic landscape in the east range of the Colombian Andes. Ninety-seven individuals belonging to eight species were collected in only two hours of sampling. The highest diversity values and the occurrence of asynanthropic flies in the forest under conservation management may reflect a positively natural restoration process in the area assessed. Grassland, rural and urban coverages were similarly low in diversity and composition. A brief discussion about the flies’ bionomy and their environment association is offered. Necrophagous flies ensembles by coverage behave as an indicator of the anthropic impact on the landscape.

## Resumen

Una evaluación ecológica rápida del ensamble de moscas necrófagas (Diptera, Calyptratae) se llevó a cabo en cuatro coberturas del paisaje antropizado del municipio de Viotá, Cundinamarca, Colombia. Noventa y siete individuos distribuidos en tres familias y ocho especies fueron recolectados en tan solo dos horas de muestreo. Los valores más altos de diversidad y la presencia de moscas asinantrópicas típicas de áreas conservadas en la cobertura de bosque, podría indicar un efecto positivo en su proceso de restauración. El pastizal, el área rural y la región urbana resultaron similares en composición y valores de baja diversidad. Se ofrece una breve discusión acerca de la bionomía de las moscas y su ambiente asociado. Finalmente evidenciamos que los ensambles de moscas por cobertura se comportan como indicadores del impacto antropogénico en el paisaje.

## Introduction

Necrophagous or carrion-feeding flies (Diptera, Calyptratae) are well known for their medico-legal, forensic and veterinary importance (Marshall, 2012); additionally, they have been suggested as a practical bioindicator of the restoration process and conservation management of the forest (de Sousa et al., 2014; Majer, 1987). Carrion-feeding flies have different tolerances to the habitat conditions, being affected by the degree of human impact on the natural environment (anthropization process) (Povolný, 1971). The preference level is commonly known in entomological studies as “Synanthropy” (Gregor & Povolný, 1958). Thus, eusynanthropic, hemisynanthropic, and asynanthropic are the ecological categories for classifying flies according to their degree of attraction or repulsion for human settlements (Povolný, 1971). In a rapid assessment (two hours), the ensemble and diversity of necrophagous flies (Diptera, Calyptratae) were assessed. Species are classified according to their associations to coverages in a heterogeneous landscape located in the east range of the Colombian Andes.

## Materials and methods

The study was conducted in a mosaic landscape area of 108 hectares, located in the Viotá Municipality, Cundinamarca province (4° 26’ 19. 73” N; 74° 31’ 12. 47” W) at 600 m of elevation in the east range of the Andean cordillera. The climate in Viotá is usually hot and humid; monthly temperature averages range from 21°C to 32°C. Four coverages of the landscape were surveyed (Forest 4° 26› 34. 15» N, 74° 30’ 47. 41”W; Grassland 4° 26› 28.93 »N, 74° 31’0. 48” O; Rural 4°26›27.28»N, 74°31›15.51»W and Urban 4° 26› 14.78 »N, 74°31’15.78”O), one collection site was settled in each (Figure 1). The secondary forest coverage is located within the Camino Verde Natural Reserve (CVNR) (Figure 1). This natural sanctuary has 100.000 m2 (10 ha).; it was traditionally used for cattle raising decades ago, but nowadays, it is under active restoration management, including sections restored from 9 to 24 years old. The grassland of *Cenchrus clandestinus* (Kikuyu grass), the rural vicinities of ranch houses, and the adjacent urbanized area of the Viotá municipality complete the set of coverages. Flies collection took place beginning the dry season on the 7th of December of 2018, starting approximately 10:00 am to 2:00 pm (two effective hours of sampling). In each assessed site, an active manual collection for thirty minutes was performed, using an entomological net, and as a bait, a three-day-old fish head rotten (sampling effort unit). Flies gathered were preserved in ethanol (C2H5OH) 90% and taxonomically identified following the keys of Amat, Vélez, and Wolff (2008); Grella et al. (2015) Wolff (2008); Grella et al. (2015); Whitworth (2014); Whitworth and Yusseff-Vanegas (2019). Pinned and labeled specimens were deposited at Colección Entomológica Tecnológico de Antioquia, Institución Universitaria (CETdeA) located in Medellín, Colombia, and also in the Colección Entomológica del Instituto Alexander Von Humboldt (IAvH) in Villa de Leiva, Colombia. The diversity data gathered by coverage was analyzed and compared based on diversity profiles and the effective number of species (Hills numbers) (Chao & Jost, 2015). Three values of diversity according to the order of coefficient *q* were plotted (the parameter *q* determines the measure of sensitivity to the species abundance); where *q*=0 for the absolute number of species (richness), *q*=1 for the exponential of Shannon entropy, and *q*=2 for the inverse of Simpson dominance. Additional Gini-Simpson equitability (*Eq*) was calculated. The category of synanthropy was deduced based on the value of the Sinanthropy Index (S.I) proposed by Nuorteva (1963) excluding individuals from grassland. The S.I ranges from −100 (avoidance for human settlements) to 100 (affinity for human settlements). Finally, synanthropic data is contrasted with previous information reported in the literature; a brief discussion about the flies’ synanthropy and their landscape associated (Figure 2) is offered.

**Figure 1.**
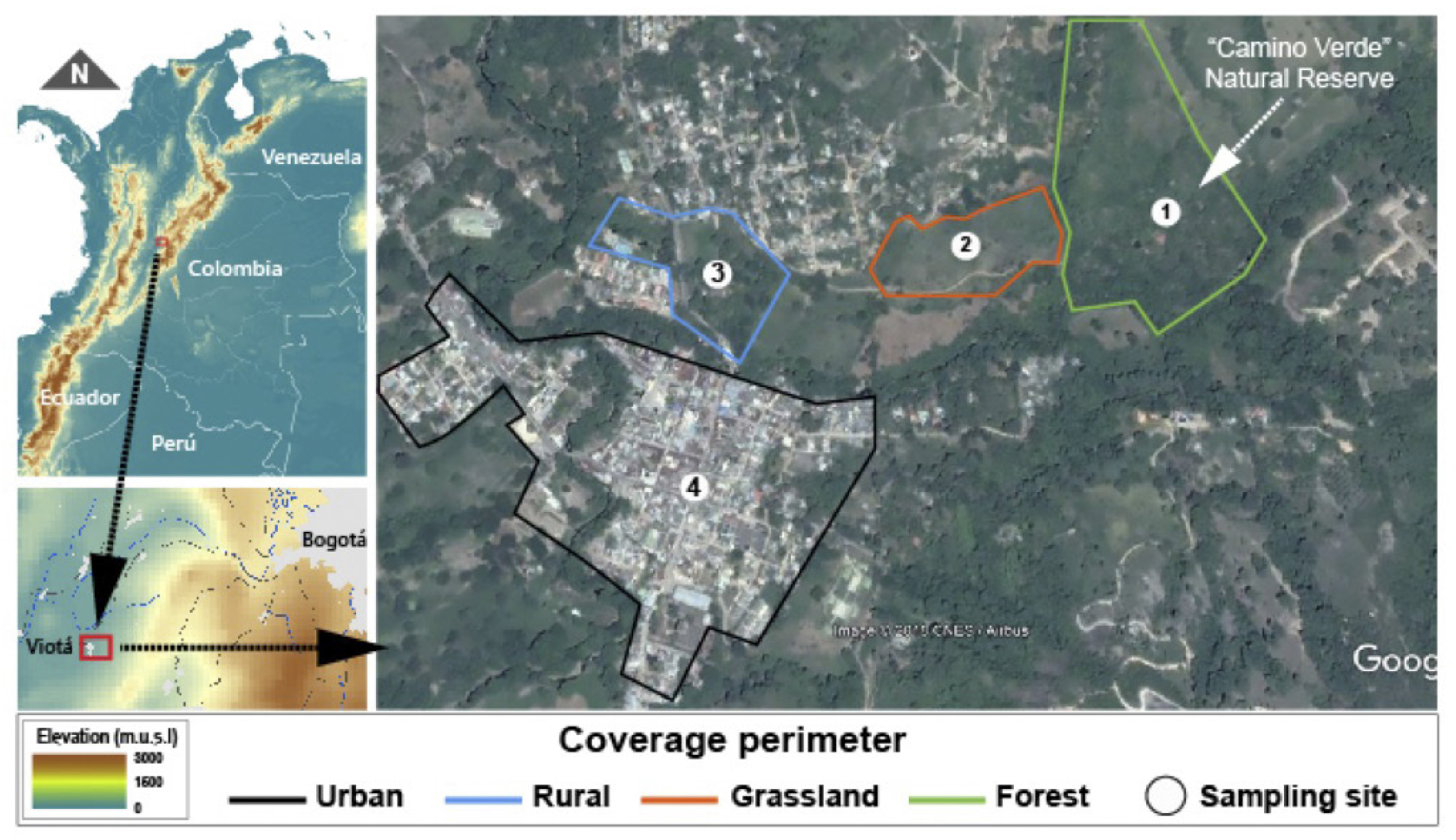
The geographical location of the study area, coverages assessed, and collection sites in Viotá Municipality, Colombia. (Satellite image extracted from Google Earth)

**Figure 2.**
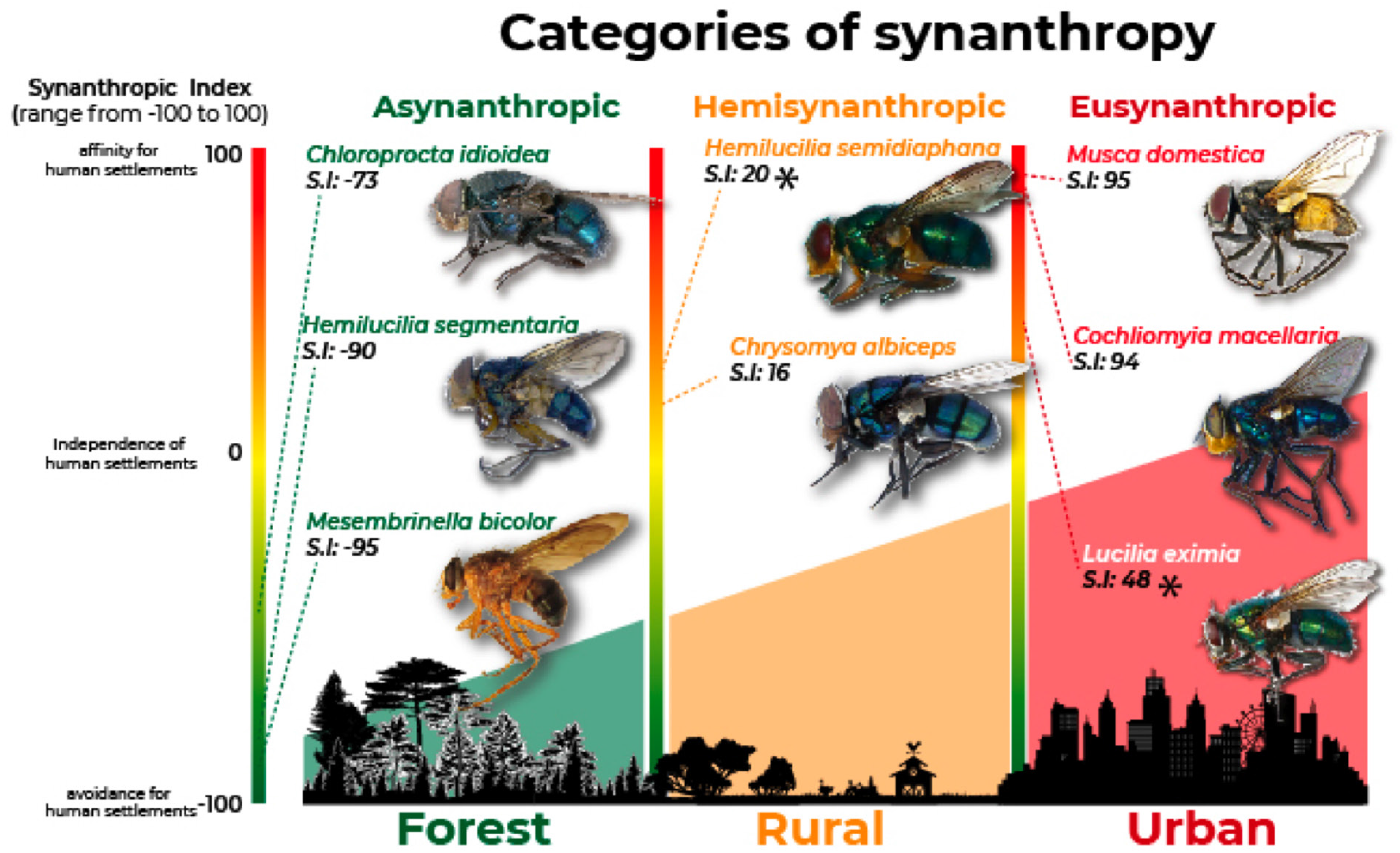
Synanthropic tendencies and categorization of necrophagous flies in the Viotá Municipality. The values of the Synanthropy Index (S.I) were based on the literature. *asynanthropic in this study.

## Results

Ninety-seven individuals (80 females, 82,4% and 17 males, 17.5%) belonging to eight species (Figure 2), encompassing three families (Calliphoridae, Mesembrinellidae, and Muscidae) of Calyptratae flies were collected. The most common species was *Chrysomya albiceps* (Wiedemann, 1819), followed by *Musca domestica* Linnaeus, 1758, and *Cochliomyia macellaria* (Fabricius, 1775). Only one individual of *Chloroprocta idioidea* (Robineau-Desvoidy, 1830), *Lucilia eximia* (Wiedemann, 1830), and *Mesembrinella bicolor* (Fabricius, 1893) were recorded (Table 1).

**Table 1.**
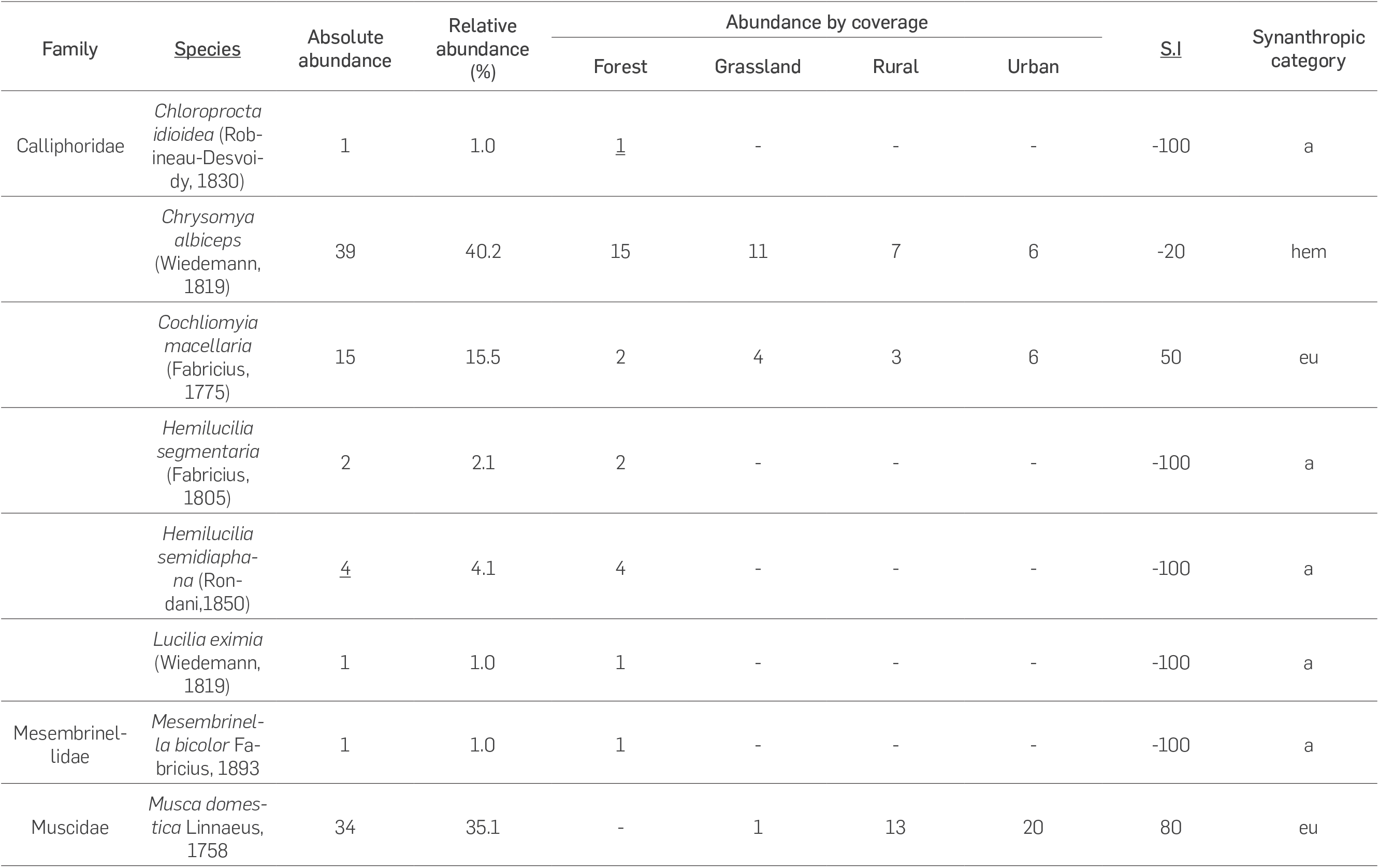
Necrophagous flies of Viotá municipality were collected in two hours sampling; abundance tendencies; Synanthropy index (S.I) and their synanthropic categorization. (a) asynanthropic; (eu) eusynanthropic; (hem) hemisynanthropic.

Necrophagous flies were most abundant in the urban area (32 individuals), while in the grassland, only 16 individuals were collected. The forest coverage displayed the highest value of diversity when *q*=0 (7 species), while in the grassland, rural and urban coverages, it was collected three species respectively. In general the forest area was the most diverse showing the highest values of diversity orders and equitability (*q*=0:7; *q*=1:4; *q*=2:2.7 – *Eq*:73.2%), followed by the rural (*q*=0:3; *q*=1:2.6; *q*=2:2.3 – *Eq*:85.6%), urban (*q*=0:3; *q*=1:2.5; *q*=2:2.7 – *Eq*:80%) and finally the grassland coverage (*q*=0:3; *q*=1:2.2; *q*=2:1.9 – *Eq*:69%) (Figure 3).

**Figure 3.**
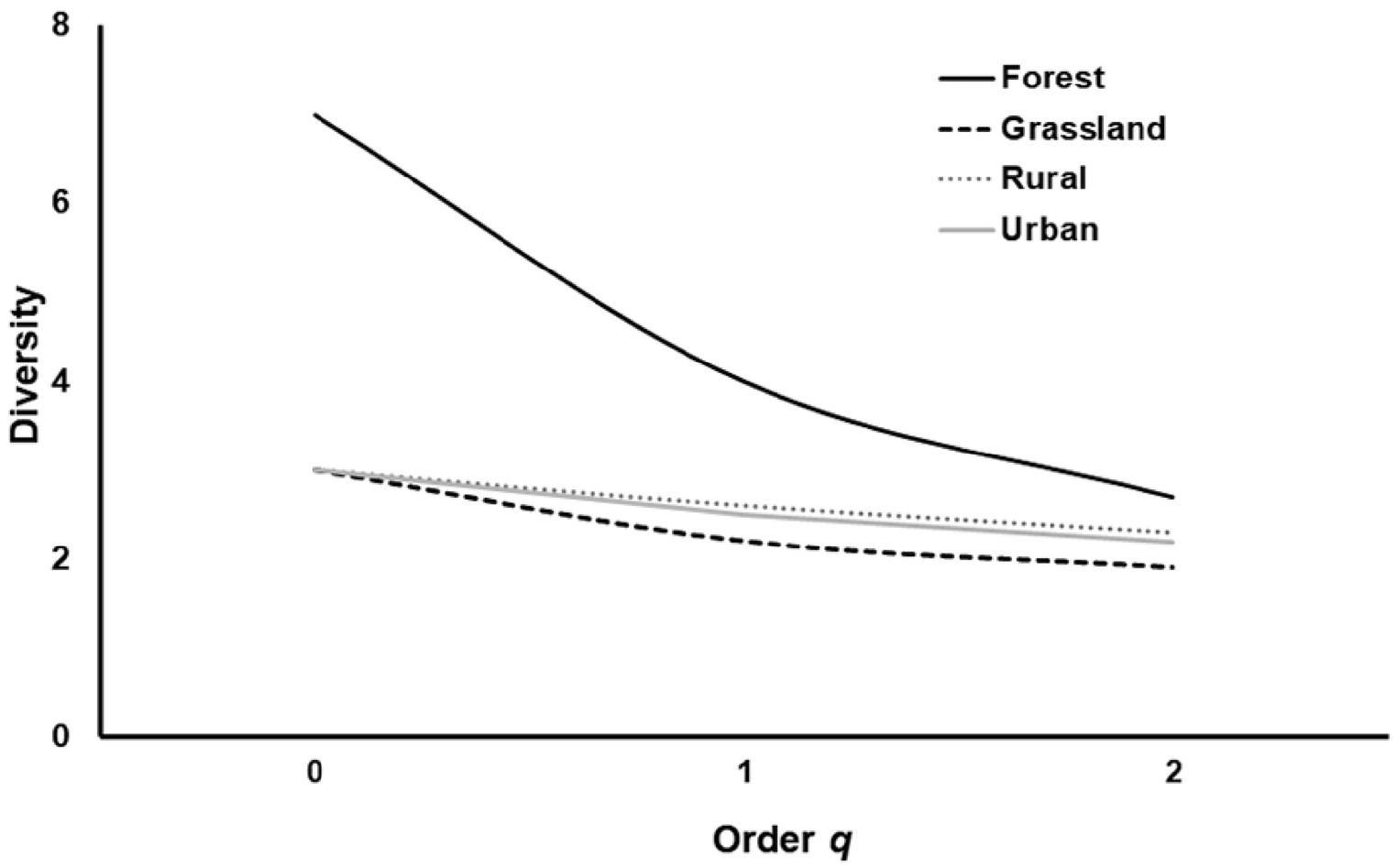
The diversity profile of the necrophagous flies based on the effective number of species (*q*=0,1 and 2) by coverage assessed in the Viotá municipality.

*M. domestica* was the most frequent species in human settlement coverages; on the other hand, *C. albiceps* was the most frequent in forest and grassland (Table 1). The forest coverage houses all the rarest, asynanthropic, and hemisynanthropic species. Synanthropic index here deduced allows to categorized five species as asynanthropic (*Chl. Idioidea*; *H. segmentaria*, *H. semidiaphana*, *L. eximia*, and *M. bicolor*); one as hemisynanthropic (*Ch. albiceps*) and finally two as eusynanthropic (*Co. macellaria*, *M. domestica*) (Table 1).

## Discussion

Despite the short time and fast sampling effort (Two hours), the richness (*q*=0) here reported (8 species) obtained in only two hours of sampling is similar with comparable Andean forensic entomological studies of flies associated with decomposed corpses during 144 hours in the fresh and bloated stage (9 species) (Grisales et al., 2010) and during 244 hours in the central range of Colombia (10 species) (Montoya-G et al., 2009). Coverage diversity data here obtained are in agreement with the study of de Sousa et al. (2020) in Amazonia, where they report the highest values of diversity estimated in the preserved environments; they also pointed that necrophagous flies respond differentially to the anthropic activity especially to cattle ranching. In other words, a more preserve environment displays higher values of diversity, and inversely the more urbanized or disturbed environments exhibit agthe lowest values. The ensemble here assessed is dominated by *C. albiceps*, a highly competitive introduced species (Rosa et al., 2006). The high abundance values of introduced species (*Chrysomya* spp.) is a common tendency in tropical carrion-flies ensembles (Amat 2017). Nevertheless, the ecological effects regarding biomass removal and competition are still unknown. According to the coverage, flies’ synanthropy previously pointed by earlier studies and the diversity tendencies agree with the landscape of Viotá (Figure 3). This means that synanthropic species were most abundant in urbanized or highly perturbed environments but scarce in the forest or better-preserved habitats and vice-versa.

All fly species in Viotá kept the ecological category previously reported in the literature except *H. semidiaphana* and *Lucilia eximia* here considered as asynanthropic. Bionomical data of *C. albiceps*, *Co macellaria*, and *Chl. Idioidea* agree with Baumgartner and Greenberg (1985) and Montoya-G, Sanchez, and Wolff (2009); *M. bicolor* with D’almeida and Lopes (1983) and *M. domestica* with Uribe-M, Wolff, and Carvalho (2010) (Figure 3). Conversely, *H. semidiaphana* was reported as hemisynanthropic in several environments of Northern South America (Amat 2017) and *L. eximia* as eusynanthropic in Amat (2017); and Montoya-G, Sanchez, and Wolff (2009). The short time sampling noticeably affects the values of abundance entered to calculate the synanthropic index. Nevertheless, for the rest of the species values here offered agree in terms of ecological classification. *H. semidiaphana* and *L. eximia* are highly polymorphic species widespread in the neotropical region (Whitwhorth 2014, Amat 2017). Therefore, there is a high probability that species complexes morphologically identical with different bionomic attributes exist, here lies the importance of local and regional ecological surveys.

The occurrence of *M. bicolor* and the other highly asynanthropic species (Figure 3) in the forest coverage suggests that it is capable of supporting their natural populations, which are particularly feasible in highly preserved and forested environments (Gadelha, 2009). However, the abundance is still low comparing to pristine environments and obviously by the sampling effort. The diversity data here obtained may foresee a preliminary positive effect of the natural process of restoration in the CVRN, indicating an adequate management, and right decisions doing livestock reconversion recovering the ecosystem services of the patch forest, in this area dominated by urbanization and cattle ranching. Although the data on adequate sampling for this ensemble may require more effort, this approach in only two hours of fieldwork, with a minimum budget and yielding this type of quality data, is undoubtedly a remarkable option to include these organisms in rapid ecological assessments (REA). The necrophagous flies assessed meets the criteria to be an optimal bioindicator of anthropic effects during environmental management processes.

Flies are abundant, easy to sample, and respond rapidly to a change in environment (synanthropy); additionally, they are relatively well taxonomic and ecological understood as here evidenced. Additional surveys conducted in other neotropical environments with different degrees of anthropic effects would corroborate flies’ efficiency as indicator taxa. For a more accurate interpretation of the synanthropy indices following this methodology, we suggest comparing values obtained with data previously obtained in ecological studies made with a more significant sampling effort to corroborate synanthropic trends among assessed populations. Including carrion-flies in rapid ecological assessments provides optimal preliminary data for local forensic entomology use and indirectly monitors landscape changes in the conservation context.

## Acknowledgments

We thank Ricardo Morales and Miguel Reyes for supporting the logistics of this survey in Reserva Natural Camino Verde at Viotá, Cundinamarca. Thanks to the staff of the Bioforense research group at Tecnológico de Antioquia, Institución Universitaria.

